# Continuous Nuclear Export of p62/SQSTM1 Is Essential for Kidney Homeostasis

**DOI:** 10.64898/2026.06.29.734648

**Authors:** Baoshuo Ning, Kunio Kawanishi, Dedong Kang, Rinna Tatsuno, Toshiaki Usui, Naoki Morito, Toru Yanagawa, Seiya Mizuno, Satoru Takahashi, Eiji Warabi

## Abstract

The selective autophagy receptor p62/SQSTM1 dynamically shuttles between the nucleus and cytoplasm through distinct nuclear localization and export signals, yet the physiological significance of this trafficking has remained unknown. Here, we generated mice carrying a deletion of the p62 nuclear export signal (dNES) to determine the in vivo role of p62 nuclear export. Homozygous dNES mice developed progressive podocyte injury, glomerulosclerosis, and fatal renal failure by 6-7 weeks of age, whereas heterozygous and dNES/- mice did not develop renal dysfunction. Loss of nuclear export caused constitutive nuclear accumulation of p62, accompanied by the formation of insoluble ubiquitin-positive aggregates and widespread alterations in the renal proteome, including activation of energy metabolism-related pathways and suppression of developmental programs. We previously demonstrated that the lipid peroxidation product 4-hydroxy-2-nonenal (4-HNE) inhibits the nuclear export receptor XPO1, resulting in nuclear retention of p62 in cultured cells. The present findings provide in vivo evidence that continuous nuclear export of p62 is indispensable for maintaining kidney homeostasis and reveal that excessive nuclear accumulation, rather than cytoplasmic depletion, underlies p62-mediated toxicity. Collectively, these findings establish continuous nuclear export of p62 as an essential mechanism for maintaining kidney homeostasis.

**Significance Statement:** The adaptor protein p62/SQSTM1 continuously shuttles between the nucleus and cytoplasm, but the physiological significance of this trafficking has remained unknown. Here, we show that disrupting the nuclear export signal of p62 causes progressive podocyte injury, glomerulosclerosis, and fatal kidney failure through excessive nuclear accumulation and aggregate formation. In contrast, dNES/+ and dNES/- mice remain healthy, demonstrating that excessive nuclear accumulation, rather than cytoplasmic depletion, drives disease. These findings identify continuous nuclear export as a fundamental mechanism that prevents toxic nuclear accumulation of p62 and preserves kidney homeostasis.

## Introduction

Many proteins continuously shuttle between the nucleus and cytoplasm through coordinated nuclear import and export. Although this dynamic trafficking is a fundamental feature of eukaryotic cells, why proteins continuously undergo this process and whether it is required for normal tissue homeostasis remain largely unknown. The selective autophagy receptor p62/SQSTM1 is one such shuttling protein. While p62 is best known for its roles in selective autophagy, ubiquitin signaling, and oxidative stress responses (1–5), it also undergoes continuous nucleocytoplasmic trafficking mediated by distinct nuclear localization signals (NLSs) and a nuclear export signal (NES) (6,7). Whether this dynamic trafficking is physiologically required has not been addressed in vivo.

p62 is ubiquitously expressed in mammalian tissues and is predominantly localized in the cytoplasm under steady-state conditions (2,3). In the cytoplasm, p62 functions as an autophagy receptor by linking ubiquitinated cargo to LC3 (4) and activates the Keap1–Nrf2 pathway through sequestration of Keap1 (5). However, inhibition of Exportin-1 (XPO1/CRM1)-dependent nuclear export by leptomycin B (LMB) rapidly induces nuclear accumulation of p62 (6,7), demonstrating that p62 continuously shuttles between the nucleus and cytoplasm despite its predominantly cytoplasmic localization. Increasing evidence further suggests that the biological functions of p62 depend not only on its abundance but also on its intracellular localization. Indeed, altered subcellular distribution of p62 has been associated with several human malignancies, including oral squamous cell carcinoma and prostate cancer, where nuclear or cytoplasmic accumulation correlates with disease progression and patient prognosis (7,8).

Mechanistically, Pankiv et al. identified two NLSs and one NES within p62 and demonstrated that these motifs mediate its nucleocytoplasmic trafficking (7). Under conditions of impaired nuclear export, p62 accumulates in promyelocytic leukemia (PML) bodies together with ubiquitinated proteins and facilitates proteasomal degradation of ataxin-1 (7). More recently, Fu et al. showed that p62 undergoes liquid–liquid phase separation (LLPS) in the nucleus, forming condensates enriched in ubiquitin-conjugating enzymes, deubiquitinating enzymes, and other components of the ubiquitin–proteasome system (UPS), thereby promoting nuclear protein quality control (6). Collectively, these studies suggest that nucleocytoplasmic trafficking enables p62 to participate in nuclear protein quality control. However, all of these observations have been limited to cultured cells, and the physiological significance of p62 nucleocytoplasmic trafficking in vivo has remained completely unknown.

To address this question, we generated mice carrying a deletion of the p62 nuclear export signal (dNES). Using this genetic model, we demonstrate that continuous nuclear export of p62 is indispensable for preventing its excessive nuclear accumulation and for maintaining kidney homeostasis. Loss of nuclear export results in nuclear p62 aggregation, progressive podocyte injury, glomerulosclerosis, and ultimately fatal renal failure, providing the first in vivo evidence that continuous nucleocytoplasmic trafficking of p62 is essential for tissue homeostasis.

## Results

### Lethal phenotype and mortality in dNES homozygous mice at 6–7 weeks of age

To investigate the physiological significance of p62 nucleocytoplasmic shuttling, we generated mutant mice harboring deletions of NES by CRISPR-Cas9 system (Fig. 1A). To examine p62 subcellular localization changes caused by NES deletion, we prepared mouse embryonic fibroblasts (MEFs). In wild-type (WT), treatment with LMB caused a clear shift in p62 localization from the cytoplasm to the nucleus, whereas dNES-p62 was nuclear even in the absence of LMB (Fig. 1B). These findings confirm that nuclear export is abolished by NES deletion.

**Figure 1.**
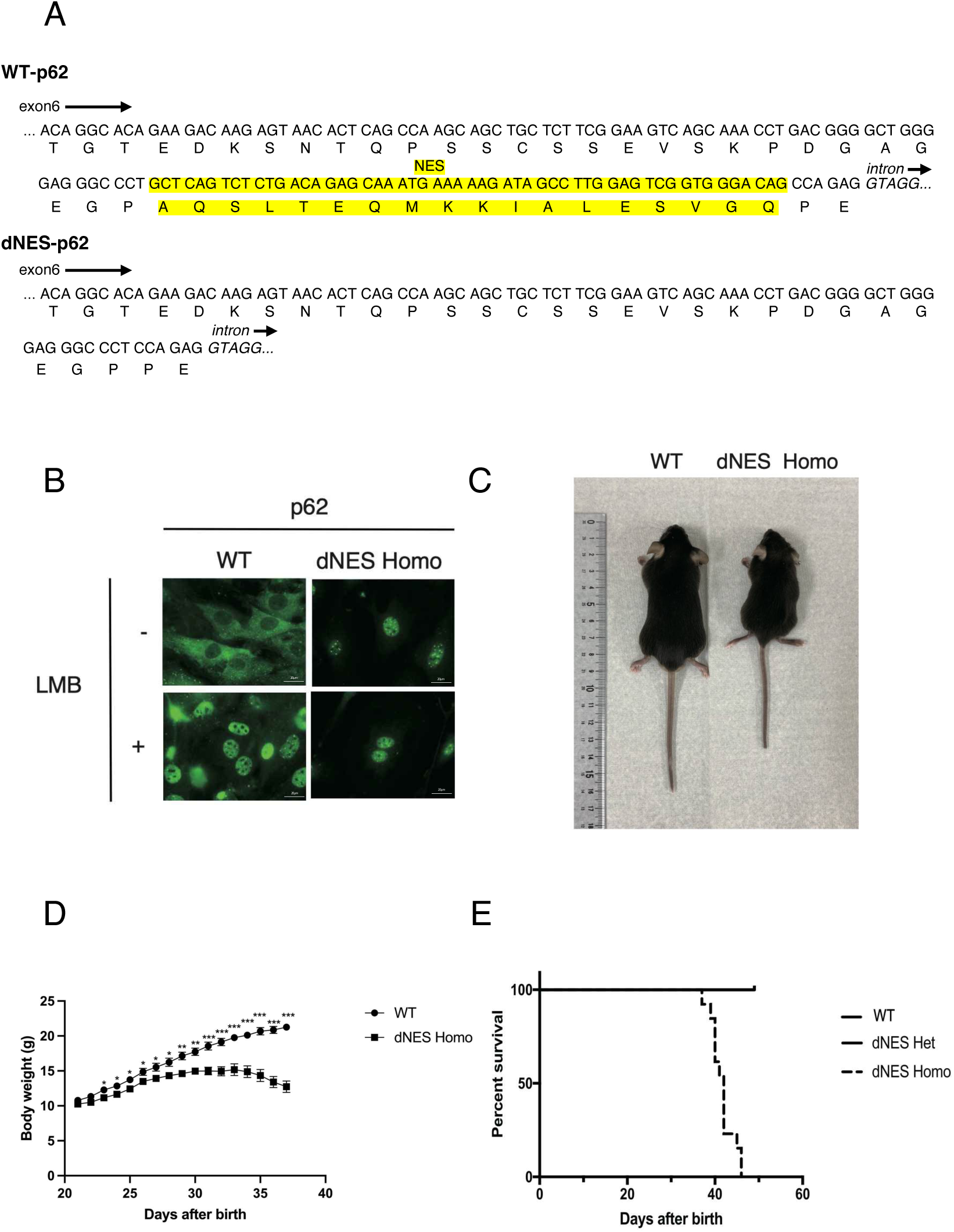
NES deletion abolishes nuclear export of p62 and causes growth retardation and early mortality in mice. (A) Schematic representation of the Sqstm1 genomic region showing the nuclear export signal (NES) highlighted in yellow. The dNES-p62 allele generated by CRISPR–Cas9 harbors an in-frame deletion that removes the NES-coding sequence. (B) Immunofluorescence analysis of p62 in MEFs. In WT MEFs, leptomycin B (LMB; 10 nM, 3 h) induced nuclear accumulation of p62, whereas dNES-p62 was constitutively nuclear even without LMB, confirming loss of nuclear export activity. Scale bars: 20 µm. (C) Representative images of WT and dNES homozygous mice at 6 weeks showing marked growth retardation. (D) Body weight curves of WT and dNES homozygous mice (WT, n = 9; homozygous, n = 10). Data are mean ± SEM. Statistical significance was determined by unpaired two-tailed Student’s t-test. *P < 0.05; **P < 0.01; ***P < 0.001. (E) Kaplan–Meier survival analysis showing that dNES homozygous mice die by 6–7 weeks of age (WT, n = 12; heterozygous, n = 12; homozygous, n = 13).

dNES homozygous mice showed growth retardation, with a smaller body size and a marked decline in body weight, and died by approximately 6–7 weeks (Fig. 1C–E). These findings demonstrate that continuous nuclear export is indispensable for maintaining normal p62 localization in vivo and that disruption of this process results in an early lethal phenotype.

### Renal dysfunction in dNES homozygous mice

Gross examination revealed that kidneys from dNES homozygous mice were paler and firmer than those of WT mice (Fig. 2A). Histological analysis by hematoxylin and eosin (HE) and periodic acid–Schiff (PAS) staining showed progressive structural abnormalities, including abundant protein casts (Fig. 2B). Quantitative evaluation of PAS-stained sections demonstrated a significant increase in focal segmental glomerulosclerosis (FSGS), crescent formation, and global sclerosis in 6-week-old dNES homozygous mice, whereas 3- and 4-week-old mice exhibited only mild alterations (Fig. 2C).

**Figure 2.**
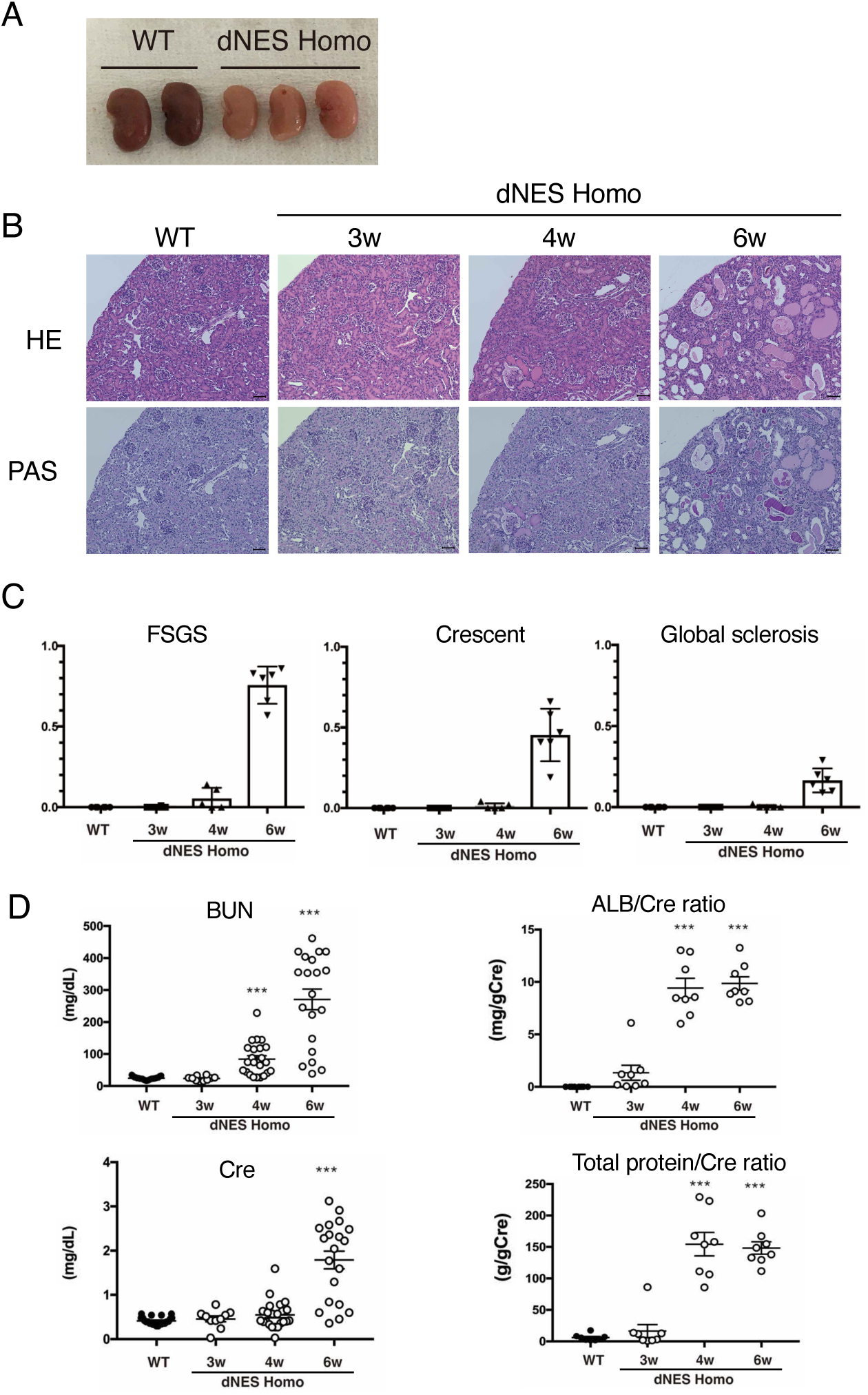
Progressive renal dysfunction and glomerular injury in dNES homozygous mice. (A) Gross morphology of kidneys from WT and dNES homozygous mice at 6 weeks. dNES kidneys are paler and firmer. (B) Representative HE and PAS staining showing glomerular and tubular abnormalities, including abundant protein casts in dNES kidneys. Scale bars: 100 µm. (C) Quantification of glomerular lesions, including focal segmental glomerulosclerosis (FSGS), crescent formation, and global sclerosis. A renal pathologist evaluated 100 glomeruli per mouse on PAS-stained sections. WT, n = 6; dNES (3 weeks), n = 6; dNES (4 weeks), n = 5; dNES (6 weeks), n = 6. Data are mean ± SEM. One-way ANOVA with Tukey’s test; ***, P < 0.001. (D) Serum biochemical measurements (BUN, creatinine, albumin/creatinine ratio, and total protein/creatinine ratio) from WT and dNES homozygous mice at 3–6 weeks. Elevated values indicate renal dysfunction beginning at ∼4 weeks. Data are mean ± SEM (one-way ANOVA with Tukey’s test; ***, P < 0.001).

Serum biochemical analysis revealed elevated creatinine (Cre) levels starting at 6 weeks, while blood urea nitrogen BUN, albumin/Cre ratio, and total protein/Cre ratio increased from 4 weeks onward (Fig. 2D). These findings indicate progressive renal dysfunction,prompting us to investigate the earliest cellular events.

### Podocyte damage detectable from 3 weeks of age

Podocyte numbers were assessed by immunostaining with the podocyte marker p57(9). The number of p57-positive cells per glomerulus declined with age in dNES homozygous mice and was significantly reduced at 6 weeks (Fig. 3A, B). Nephrin immunofluorescence, reflecting slit diaphragm integrity, also decreased progressively and was nearly undetectable at 6 weeks (Fig. 3C). Western blot analysis confirmed a marked reduction in Nephrin protein in 6-week-old dNES homozygous mice (Fig. 3D).

**Figure 3.**
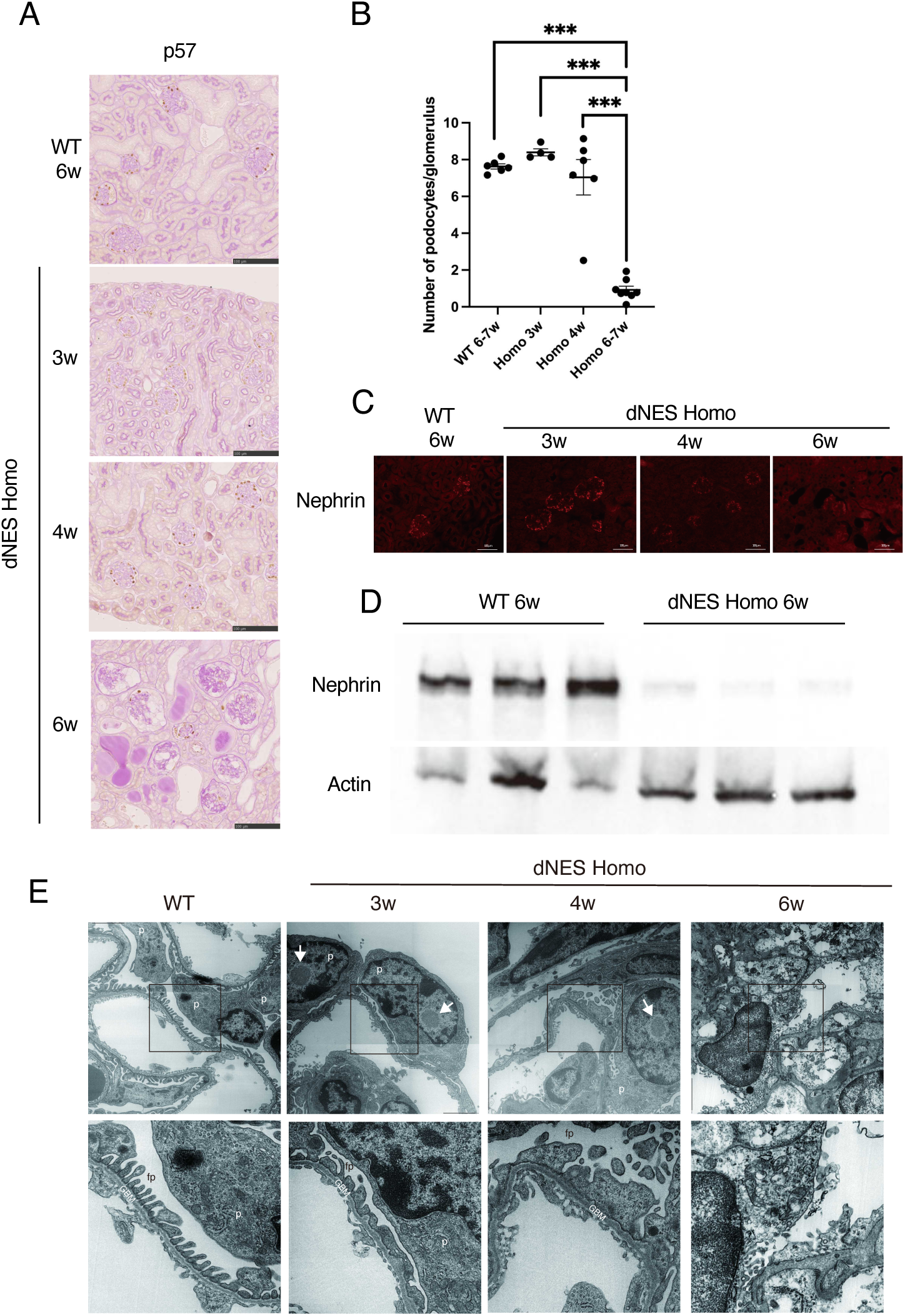
Early and progressive podocyte injury in dNES homozygous mice. (A) Immunohistochemical staining for the podocyte marker p57 showing an age-dependent decrease in p57-positive cells in dNES kidneys. Scale bars: 100 µm. (B) Quantification of podocyte number per glomerulus. For each mouse, 100 glomeruli were analyzed and p57-positive nuclei counted. WT (6–7 weeks): n = 6; dNES (3 weeks): n = 4; dNES (4 weeks): n = 6; dNES (6 weeks): n = 8. Data are mean ± SEM (one-way ANOVA with Tukey’s test; ***, P < 0.001). (C) Immunofluorescence staining for Nephrin, showing progressive loss of slit diaphragm signal by 6 weeks. Scale bars: 100 µm. (D) Western blot analysis of Nephrin in kidney lysates, with Actin as a loading control. (E) Transmission electron microscopy showing foot process effacement (black boxes) and electron-dense aggregates (white arrows) in podocytes of dNES mice as early as 3 weeks. p: podocyte; fp: foot process; GBM: glomerular basement membrane.

Transmission electron microscopy revealed podocyte foot process effacement as early as 3 weeks, accompanied by unidentified electron-dense aggregates (Fig. 3E). These observations indicate that podocyte injury precedes overt renal failure, suggesting that podocytes are the primary cellular target of impaired p62 nuclear export.

### Nuclear co-localization of p62 and ubiquitin in the kidneys of dNES homozygous mice

Immunostaining of kidney sections showed nuclear p62-positive puncta in dNES homozygous mice at 4 and 6 weeks, which were absent in WT controls (Fig. 4A). Co-immunostaining revealed that ubiquitin colocalized with p62 within the nucleus, suggesting accumulation of ubiquitinated aggregates.

**Figure 4.**
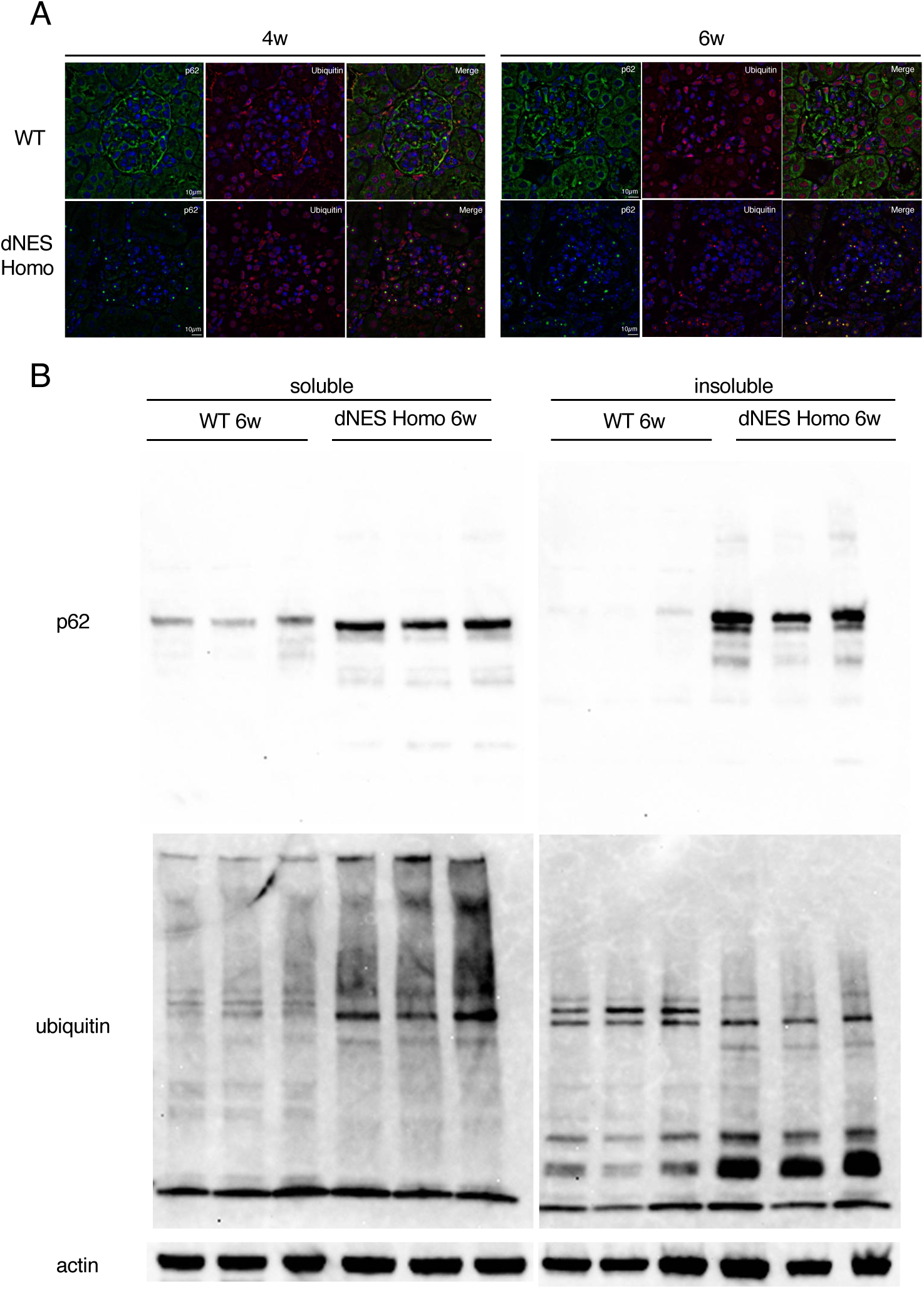
Nuclear accumulation of p62–ubiquitin aggregates in dNES homozygous mice. (A) Immunofluorescence staining of kidney sections showing cytoplasmic p62 in WT kidneys and nuclear p62 puncta co-localized with ubiquitin in dNES kidneys at 4 and 6 weeks. Nuclei are counterstained with Hoechst. (B) Western blot analysis of soluble and insoluble kidney fractions. p62 and ubiquitin are markedly enriched in the insoluble fraction of dNES kidneys, consistent with aggregate formation.

Western blot analysis of soluble and insoluble fractions demonstrated that p62 was enriched in the insoluble fraction of dNES homozygous kidneys (Fig. 4B), consistent with aggregate formation. Ubiquitin levels were also elevated in both soluble and insoluble fractions, indicating a broad impairment of ubiquitinated protein clearance. The increase in soluble p62 and ubiquitinated proteins suggests that not only aggregate-prone species but also normally degradable substrates accumulate, reflecting a global disruption of p62-dependent protein quality control. Together, these findings indicate that failure of p62 nuclear export disrupts nuclear protein quality control, leading to accumulation of both insoluble aggregates and soluble ubiquitinated proteins.

### Excessive nuclear accumulation of p62 underlies kidney injury

To determine whether kidney injury results simply from loss of nuclear export or from the extent of nuclear p62 accumulation, we generated dNES/– mice by crossing dNES heterozygotes with p62-knockout mice. Despite expressing export-defective p62, dNES/– mice survived normally and exhibited body weights and BUN levels comparable to those of dNES/+ mice (Fig. 5A). Immunofluorescence analysis showed abundant nuclear p62 aggregates in dNES homozygous kidneys, whereas dNES/+ and dNES/- mice displayed only a few nuclear p62-positive puncta (Fig. 5B). Consistently, western blot analysis demonstrated that total p62 expression in dNES/- kidneys was comparable to that in WT mice and substantially lower than that in dNES homozygotes (Fig. 5C). Together, these findings demonstrate that kidney injury is determined by the extent of nuclear p62 accumulation rather than by the mere loss of nuclear export, suggesting that excessive nuclear accumulation of p62 beyond a critical level drives renal pathology.

**Figure 5.**
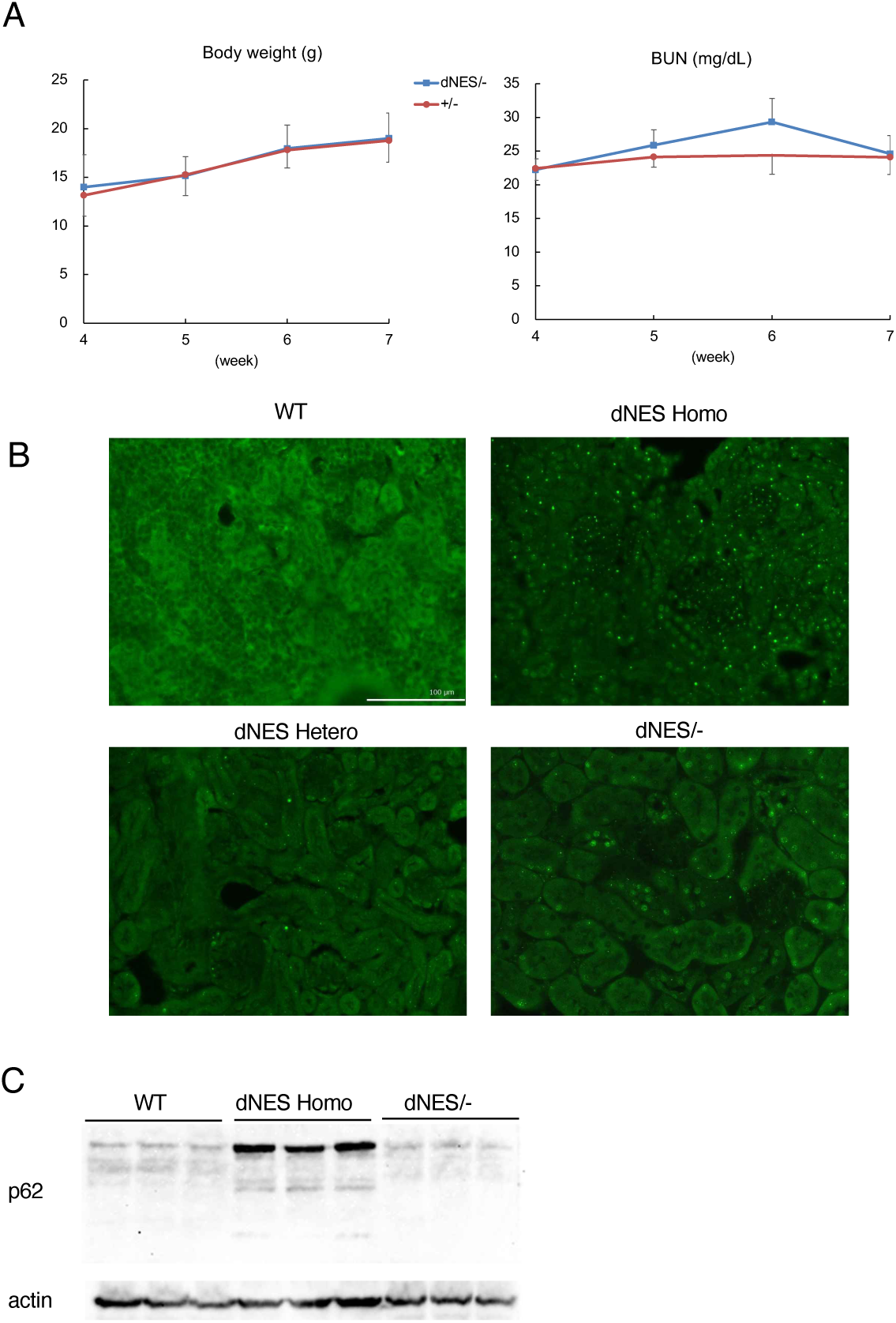
Reduced p62 dosage limits nuclear aggregation and prevents renal failure in dNES/– mice. (A) Body weight and blood urea nitrogen (BUN) levels of dNES heterozygous and dNES/– mice showing normal growth and kidney function. (B) Immunofluorescence staining for p62 in kidneys from WT, dNES heterozygous, dNES homozygous, and dNES/– mice. Abundant nuclear aggregates are present in homozygotes but reduced in heterozygotes and dNES/– mice. (C) Western blot analysis of p62 expression in kidney lysates prepared with RIPA buffer.

### Proteomic analysis reveals dysregulation of protein expression profiles

Principal component analysis (PCA) of proteomic data clearly separated WT and dNES homozygous samples along PC1 (36%) and PC2 (20%) axes, indicating major proteomic differences (Fig. 6A). A volcano plot highlighted numerous significantly up- and down-regulated proteins in dNES homozygous mice (Fig. 6B). Quantitative comparison showed more up-regulated than down-regulated proteins (Fig. 6C). Heatmap clustering revealed distinct expression patterns between the two groups (Fig. 6D). Gene Ontology enrichment analysis identified upregulated pathways related to energy metabolism and downregulated pathways associated with cell and tissue development (Fig. 6E). Together, these analyses demonstrate that loss of p62 nuclear export profoundly alters renal proteostasis, with altered metabolic pathways and impaired developmental and structural maintenance programs.

**Figure 6.**
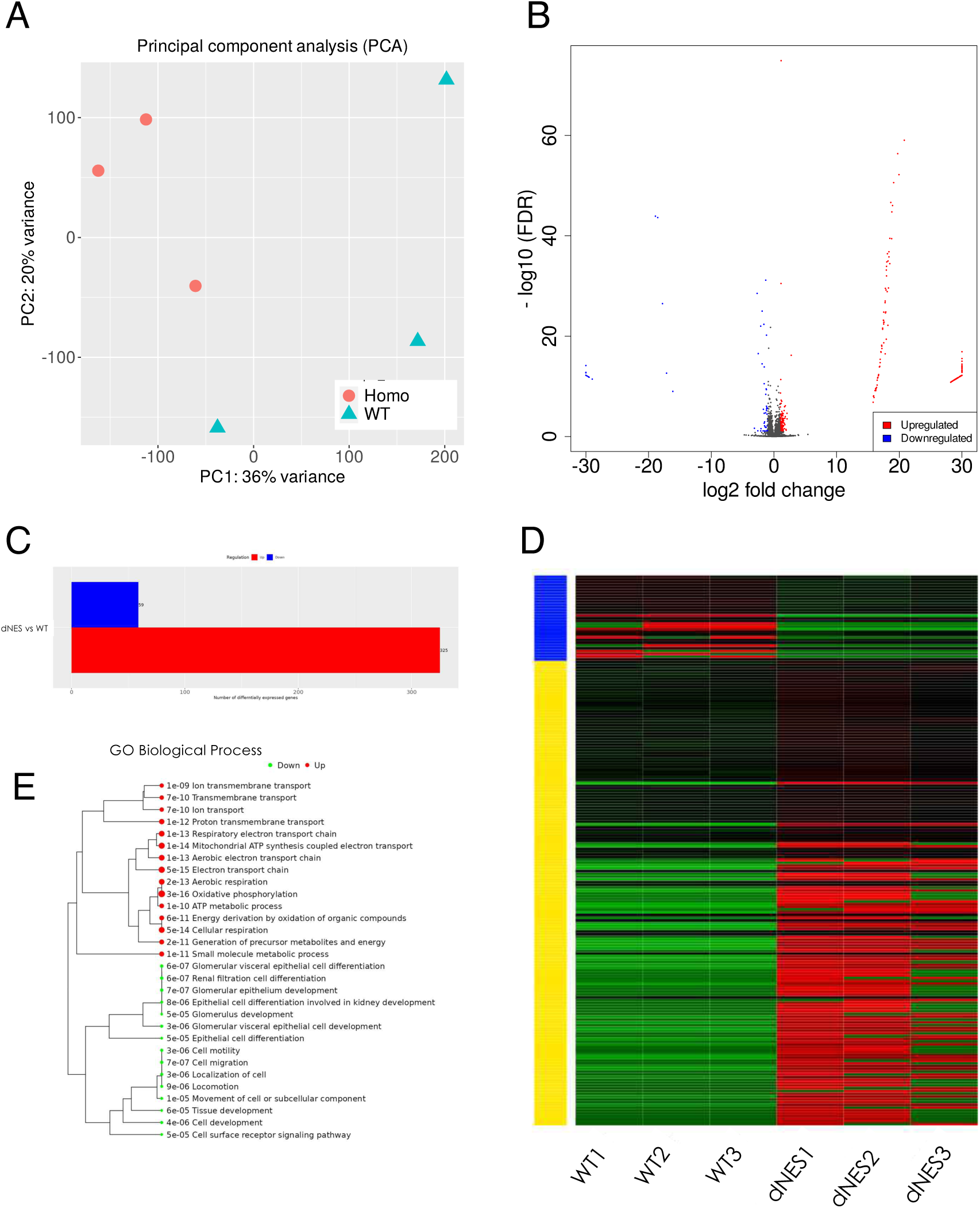
Proteomic profiling reveals widespread changes in protein abundance and metabolic reprogramming in dNES kidneys. (A) Principal component analysis (PCA) showing clear separation of WT and dNES homozygous kidney samples. (B) Volcano plot showing significantly upregulated (red) and downregulated (blue) proteins in dNES kidneys. (C) Summary of the number of upregulated versus downregulated proteins. (D) Heatmap with hierarchical clustering showing distinct proteomic signatures between WT and dNES samples. (E) Gene Ontology (GO) enrichment analysis of upregulated (red) and downregulated (green) protein categories, highlighting increased energy metabolism and reduced developmental pathways in dNES kidneys.

## Discussion

The present study provides the first in vivo evidence that continuous nuclear export of p62 is indispensable for maintaining kidney homeostasis. By disrupting the nuclear export signal of p62, we demonstrated that constitutive nuclear retention of p62 leads to nuclear accumulation of ubiquitin-positive aggregates, progressive podocyte injury, glomerulosclerosis, and ultimately fatal renal failure. These findings establish that continuous nuclear export is not simply a mechanism for intracellular protein transport but is required to maintain normal p62 localization and renal function.

An important finding of this study is that kidney injury was associated with the extent of nuclear p62 accumulation rather than with the mere loss of nuclear export itself. Although both dNES homozygous and dNES/– mice expressed export-defective p62, only dNES homozygotes developed renal disease. In contrast, dNES/– mice, which expressed substantially lower amounts of mutant p62, exhibited only limited nuclear accumulation and remained free of renal dysfunction. These observations suggest that excessive nuclear accumulation of p62 beyond a critical level, rather than simple impairment of nuclear export, is a key determinant of renal pathology.

Previous studies have primarily focused on the cytoplasmic functions of p62 as a selective autophagy receptor and signaling scaffold regulating pathways such as Keap1–Nrf2 signaling (5,13,14). In contrast, the physiological significance of nuclear p62 has remained largely unexplored. Cell culture studies demonstrated that p62 localizes to PML bodies under conditions of nuclear export inhibition and participates in ubiquitin–proteasome system (UPS)-dependent degradation of nuclear proteins (6,7). More recently, p62 was shown to undergo liquid–liquid phase separation (LLPS) within the nucleus, generating condensates enriched in ubiquitin-conjugating enzymes, deubiquitinating enzymes, and other UPS components (6). Our in vivo findings extend these observations by demonstrating that continuous nuclear export is required to prevent excessive nuclear accumulation of p62 and preserve organ function. The accumulation of both insoluble p62–ubiquitin aggregates and soluble ubiquitinated proteins in dNES kidneys further suggests that disruption of p62 trafficking impairs nuclear protein quality control rather than simply promoting aggregate formation.

Proteomic analysis revealed extensive remodeling of the renal proteome, characterized by upregulation of metabolism-related pathways and downregulation of developmental programs. Although the causal relationship between these proteomic changes and nuclear p62 accumulation remains to be established, these findings are consistent with chronic cellular stress accompanied by progressive deterioration of tissue homeostasis. Future studies combining time-resolved proteomics with single-cell transcriptomic analyses will help define the molecular events linking p62 nuclear retention to podocyte dysfunction.

The mechanisms by which nuclear p62 accumulation causes renal injury remain to be determined. Several, non-mutually exclusive possibilities may be considered. Excessive nuclear p62 condensates may sequester ubiquitin and components of the UPS, thereby impairing nuclear protein turnover. Alternatively, aberrant condensates may interfere with the dynamics of nuclear phase-separated compartments such as PML bodies. Nuclear retention of p62 may also perturb transcriptional regulation through abnormal interactions with chromatin-associated proteins or transcriptional regulators. Elucidating these mechanisms will provide important insight into the physiological functions of nuclear p62.

An important question raised by this study is why renal pathology predominates despite ubiquitous expression of mutant p62. Podocytes are highly specialized terminally differentiated cells that depend on efficient protein quality control to maintain the glomerular filtration barrier (15). Their remarkable susceptibility to nuclear p62 accumulation is therefore consistent with previous studies showing that impairment of proteasome function or proteostasis preferentially compromises podocyte integrity and promotes glomerular disease (16–18). The foot process effacement and progressive glomerulosclerosis observed in dNES mice resemble pathological features of human focal segmental glomerulosclerosis (FSGS), suggesting that impaired p62 trafficking may represent a previously unrecognized molecular mechanism contributing to podocyte injury.

Our previous study demonstrated that the lipid peroxidation product 4-hydroxy-2-nonenal (4-HNE) inhibits XPO1, resulting in nuclear retention of p62 in cultured cells (10). Because oxidative stress and 4-HNE accumulation are common features of chronic kidney disease (11,12), the present dNES mouse provides genetic evidence that impaired nuclear export of p62 alone is sufficient to initiate progressive renal injury. These findings raise the possibility that oxidative stress-induced suppression of nuclear export contributes directly to kidney disease through abnormal nuclear accumulation of p62.

In conclusion, our study identifies continuous nuclear export of p62 as an essential mechanism for maintaining kidney homeostasis in vivo. More broadly, our findings suggest that dynamic intracellular localization represents an important layer of protein regulation in addition to protein abundance, post-translational modification, and protein turnover. We propose that continuous nucleocytoplasmic trafficking enables p62 to maintain cellular homeostasis by preventing excessive nuclear accumulation and that disruption of this dynamic equilibrium can lead to tissue-specific disease. Further investigation of the mechanisms regulating p62 nucleocytoplasmic trafficking may therefore provide new insights into the pathogenesis of protein aggregation disorders and identify novel therapeutic opportunities for kidney disease.

## Materials and Methods

### Experimental animals

The dNES mouse line was generated at the Laboratory Animal Resource Center, University of Tsukuba, using the CRISPR–Cas9 system. Two single-guide RNAs (sgRNAs) were designed to flank the NES region of the Sqstm1 gene (5′-CCTGCTCAGTCTCTGACAGAGCA-3′ and 5′-TCGGTGGGACAGCCAGAGGTAGG-3′). An ssODN donor (5′-CCCAGAAAGTTCCAGCACAGGCACAGAAGACAAGAGTAACACTCAGCCAAGCAGCTGCTCTTCGGAAGTCAGCAAACCTGACGGGGCTGGGGAGGGCCCTCCAGAGGTAGGTCTACTAGCTTCAGCCTGAGGAATCCTGTCCTTCTACTGTTACCCTGAGCCTCCTGATAGAACTCTGTAGAGAAAATTTACTTAGCTCT-3′) lacking the NES sequence was used for homology-directed repair. Initial genome editing yielded a mutant allele generated by non-homologous end joining that directly rejoined the cut sites without ssODN integration. To refine the mutation, a second round of genome editing was performed using sperm from the initial mutant line. Fertilized embryos were co-injected with Cas9 protein, sgRNA (5′-AAGCTAGTAGACCTACCAGC-3′), and an ssODN donor (5′-GGTAACAGTAGAAGGACAGGATTCCTCAGGCTGAAGCTAGTAGACCTACCTCTGGAGGGCCCTCCCCAGCCCCGTCAGGTTTGCTGACTTCCGAAGAGCAGCTGC-3′). Founders were screened by PCR using the following primers: Forward 5′-AGCACAGGCACAGAAGACAA-3′ and Reverse 5′-AGGCCTAAGCAATGACAAGT-3′, producing amplicons of 340 bp for wild-type and 286 bp for the mutant allele. The genomic sequences of mice predicted to carry the mutant allele were analyzed, confirming that the NES region was successfully deleted.

*p62*-knockout (*p62*–KO) mice were described previously (22). All mice were maintained under specific pathogen–free conditions at 25 °C with a 14-h light/10-h dark cycle and were provided standard chow and water ad libitum. All experimental protocols were reviewed and approved by the Animal Care and Use Committee of the University of Tsukuba and conducted in accordance with institutional guidelines for the care and use of laboratory animals.

### Cell culture

Mouse embryonic fibroblasts (MEFs) were prepared from wild-type (WT) and dNES homozygous embryos as previously described (23). Cells were maintained in Dulbecco’s Modified Eagle Medium (DMEM; Gibco, USA) supplemented with 10% fetal bovine serum (FBS; Gibco), 1% L-glutamine, 1% sodium pyruvate, and 1% penicillin–streptomycin at 37°C in a humidified atmosphere of 5% CO₂.

### Body weight measurement and survival analysis

Body weight was measured weekly from 3 to 6 weeks of age in WT and dNES homozygous mice. Survival was monitored daily for WT, dNES heterozygous, and dNES homozygous mice throughout their natural lifespan.

### Serum preparation and biochemical analysis

Blood samples were collected and allowed to clot at 25°C for 30 min, followed by incubation at 4°C for 16 h. Samples were centrifuged at 1,000–1,200 × g for 20–30 min at 4°C, and the supernatant serum was collected. Serum biochemical analyses were outsourced to Oriental Yeast Co., Ltd. (Tokyo, Japan).

### Immunofluorescence staining of cultured cells

Cells cultured on 35 mm dishes were fixed with 4% paraformaldehyde (Nacalai Tesque, Japan) for 30 min at room temperature, permeabilized with 0.1% Triton X-100 (Wako, Japan) in PBS for 30 min, and blocked with 1% bovine serum albumin (BSA; Nacalai Tesque) in PBS for 1 h. Cells were incubated overnight at 4°C with rabbit anti-p62 antibody (1:400) (24), followed by Alexa Fluor–conjugated secondary antibodies (1:800; Invitrogen, USA) for 1 h at room temperature. The stained cells were examined using a fluorescence microscope (BZ-X810, Keyence, Japan).

### Immunofluorescence staining of tissue sections

Both paraffin-embedded and frozen kidney sections were used. Paraffin sections were deparaffinized with xylene, rehydrated through graded ethanol, and subjected to antigen retrieval by autoclaving in Target Retrieval Solution (Dako, USA) at 121°C for 10 min. Frozen sections were equilibrated to room temperature and washed in PBS to remove the O.C.T. compound. After permeabilization with PBS containing 0.25% Triton X-100, sections were blocked with PBS containing 1% BSA and 5% goat serum, then incubated with primary and secondary antibodies as described above. Sections were mounted with Fluoromount (DBS, Netherlands) containing Hoechst 33342 and sealed with nail polish. Control sections were included in each experiment.

### Kidney sample preparation for histological analysis

WT (6–7 weeks), dNES heterozygous, dNES homozygous, and dNES/– mice (3–6 weeks) were anesthetized with isoflurane (FUJIFILM, Japan) and perfused with PBS followed by 4% paraformaldehyde (Nacalai Tesque). Kidneys were fixed in Mildform 10N (FUJIFILM) for 48 h, embedded in paraffin, or frozen in OCT compound. Paraffin sections were used for HE, PAS, and immunostaining, while frozen sections were used for immunofluorescence. HE and PAS staining were performed by the Medical Laboratory for Tissue Sample Preparation, University of Tsukuba.

### p57 immunohistochemistry and histological quantification

p57 immunostaining was performed using the manufacturer’s protocol for paraffin sections (Proteintech, USA). Focal segmental glomerulosclerosis (FSGS), crescent formation, global sclerosis, and p57-positive podocytes were quantified in PAS- and p57-stained sections, and the percentage of affected glomeruli was calculated.

### Transmission electron microscopy

Kidney tissues were fixed in 2.5% glutaraldehyde and processed for electron microscopy at the Electron Microscope Facility, Institute of Medicine, University of Tsukuba.

### Western blotting analysis

Kidney tissues were homogenized in RIPA buffer (Nacalai Tesque) supplemented with protease inhibitors. For soluble and insoluble fractionation, samples were processed as described by Komatsu *et al.* (2007). Proteins were separated on 7.5–10% SDS–PAGE gels (TGX FastCast Acrylamide Kit, Bio-Rad) and transferred to PVDF membranes using a Trans-Blot Turbo system (Bio-Rad). Membranes were blocked with Blocking One (Nacalai Tesque) and probed overnight at 4 °C with primary antibodies, followed by HRP-conjugated secondary antibodies. Signals were detected using Chemi-Lumi One Super (Nacalai Tesque) and imaged on an iBright FL1500 system (Thermo Fisher Scientific, USA).

### Proteomic analysis

Glomeruli were isolated from 4-week-old WT and dNES homozygous mice by laser capture microdissection, and proteins were analyzed by data-independent acquisition (DIA) mass spectrometry.

### Antibodies

Primary antibodies: Anti-multi-ubiquitin mAb (MBL, Japan); ubiquitin pAb (Proteintech, USA); mouse Nephrin Ab (R&D Systems, USA); Actin (C-11, sc-1615, Santa Cruz); p57 Kip (KP39, sc-56341, Santa Cruz).

Secondary antibodies: Alexa Fluor 488/546–conjugated antibodies (Invitrogen, USA); HRP-conjugated goat anti-rabbit and donkey anti-goat IgG (Invitrogen, USA).

## Data Availability

Proteomic data have been deposited in the Japan ProteOme STandard Repository (jPOST) under the accession ID JPST002023.

## Acknowledgments and funding sources

We are grateful to Dr. T. Ishii (University of Tsukuba) and Dr. G. E. Mann (King’s College London) for their critical discussions. We sincerely thank Dr. K. Asanuma (Chiba University) for valuable comments and stimulating discussions that were instrumental in advancing this research. We also thank Ms. M. Kiuchi (University of Tsukuba) for maintaining the mouse colonies, Ms. Y. Jinzenji (University of Tsukuba) for preparing electron microscopy samples, and Ms. H. Kato (University of Tsukuba) for technical assistance in generating the dNES mice. This work was supported by JSPS KAKENHI Grant Numbers 20K07421, 23K06496 (to E.W.) and 19H03846, 22H03258 (to T.Y.).

## Author Contributions

E.W. and S.T. designed the study; S.M. generated mutant mice; B.N., K.K., D.K., R.T., and E.W. conducted the study; B.N., K.K., T.U., N.M., and T.Y. analyzed the data; and E.W., B.N., and K.K. wrote the manuscript.

## References

1. A. V. Kumar, J. Mills, L. R. Lapierre, Selective Autophagy Receptor p62/SQSTM1, a Pivotal Player in Stress and Aging. Front Cell Dev Biol 10, 793328 (2022).

2. Y. Katsuragi, Y. Ichimura, M. Komatsu, p62/SQSTM1 functions as a signaling hub and an autophagy adaptor. FEBS J 282, 4672–4678 (2015).

3. S.-J. Jeong, X. Zhang, A. Rodriguez-Velez, T. D. Evans, B. Razani, p62/SQSTM1 and Selective Autophagy in Cardiometabolic Diseases. Antioxid Redox Signal 31, 458–471 (2019).

4. S. Pankiv, et al., p62/SQSTM1 binds directly to Atg8/LC3 to facilitate degradation of ubiquitinated protein aggregates by autophagy. J Biol Chem 282, 24131–24145 (2007).

5. M. Komatsu, et al., The selective autophagy substrate p62 activates the stress responsive transcription factor Nrf2 through inactivation of Keap1. Nat Cell Biol 12, 213–223 (2010).

6. A. Fu, V. Cohen-Kaplan, N. Avni, I. Livneh, A. Ciechanover, p62-containing, proteolytically active nuclear condensates, increase the efficiency of the ubiquitin-proteasome system. Proc Natl Acad Sci U S A 118, e2107321118 (2021).

7. S. Pankiv, et al., Nucleocytoplasmic shuttling of p62/SQSTM1 and its role in recruitment of nuclear polyubiquitinated proteins to promyelocytic leukemia bodies. J Biol Chem 285, 5941–5953 (2010).

8. R. Takasaki, et al., p62 Is a Potential Biomarker for Risk of Malignant Transformation of Oral Potentially Malignant Disorders (OPMDs). Curr Issues Mol Biol 45, 7630–7641 (2023).

9. M. Nagata, K. Nakayama, Y. Terada, S. Hoshi, T. Watanabe, Cell cycle regulation and differentiation in the human podocyte lineage. Am J Pathol 153, 1511–1520 (1998).

10. E. Kayama, et al., 4-Hydroxy-2-nonenal causes nuclear accumulation of p62 by inhibiting Xpo1 and promoting the proteolytic pathway in the nucleus. PLoS One 20, e0316558 (2025).

11. M. Gyurászová, R. Gurecká, J. Bábíčková, Ľ. Tóthová, Oxidative Stress in the Pathophysiology of Kidney Disease: Implications for Noninvasive Monitoring and Identification of Biomarkers. Oxid Med Cell Longev 2020, 5478708 (2020).

12. Y. Li, et al., Oxidative Stress and 4-hydroxy-2-nonenal (4-HNE): Implications in the Pathogenesis and Treatment of Aging-related Diseases. Journal of Immunology Research 2022, 2233906 (2022).

13. Y. Ichimura, et al., Phosphorylation of p62 Activates the Keap1-Nrf2 Pathway during Selective Autophagy. Molecular Cell 51, 618–631 (2013).

14. A. Jain, et al., p62/SQSTM1 is a target gene for transcription factor NRF2 and creates a positive feedback loop by inducing antioxidant response element-driven gene transcription. J Biol Chem 285, 22576–22591 (2010).

15. A. V. Cybulsky, Endoplasmic reticulum stress, the unfolded protein response and autophagy in kidney diseases. Nat Rev Nephrol 13, 681–696 (2017).

16. S.-I. Makino, et al., Impairment of Proteasome Function in Podocytes Leads to CKD. J Am Soc Nephrol 32, 597–613 (2021).

17. L. Heintz, C. Meyer-Schwesinger, The Intertwining of Autophagy and the Ubiquitin Proteasome System in Podocyte (Patho)Physiology. Cell Physiol Biochem 55, 68–95 (2021).

18. C. Delrue, M. M. Speeckaert, Renal Implications of Dysregulated Protein Homeostasis: Insights into Ubiquitin-Proteasome and Autophagy Systems. Biomolecules 15, 349 (2025).

19. W. Kriz, M. LeHir, Pathways to nephron loss starting from glomerular diseases-insights from animal models. Kidney Int 67, 404–419 (2005).

20. S. Alberti, A. A. Hyman, Biomolecular condensates at the nexus of cellular stress, protein aggregation disease and ageing. Nat Rev Mol Cell Biol 22, 196–213 (2021).

21. S. F. Banani, H. O. Lee, A. A. Hyman, M. K. Rosen, Biomolecular condensates: organizers of cellular biochemistry. Nat Rev Mol Cell Biol 18, 285–298 (2017).

22. K. Okada, et al., The alpha-glucosidase inhibitor acarbose prevents obesity and simple steatosis in sequestosome 1/A170/p62 deficient mice. Hepatol Res 39, 490–500 (2009).

23. H. Harada, et al., Deficiency of p62/Sequestosome 1 causes hyperphagia due to leptin resistance in the brain. J Neurosci 33, 14767–14777 (2013).

24. T. Ishii, et al., Transcription Factor Nrf2 Coordinately Regulates a Group of Oxidative Stress-inducible Genes in Macrophages*. Journal of Biological Chemistry 275, 16023–16029 (2000).

