## Supplemental Table 1 for "Continuous Nuclear Export of p62/SQSTM1 Is Essential for Kidney Homeostasis"

**Table S1.**

**Serum biochemical parameters in WT dNES heterozygous and homozygous mice.**

| | | AST (U/L) | ALT (U/L) |
|---|---|---|---|
| 4w | WT | 96.2±11.7 | 62.4±11.4 |
| | dNES/+ | 87.0±5.9 | 92.2±47.1 |
| | dNES/dNES | 109.9±11.9 | 39.4±3.6 |
| 7w | WT | 110.6±60.5 | 133.2±36.2 |
| | dNES/+ | 97.7±21.0 | 104.6±65.2 |
| | dNES/dNES | 136.0±20.0 | 37.2±6.4 |

AST and ALT levels were measured in WT, dNES heterozygous, and dNES homozygous mice at 4 and 7 weeks of age. Although ALT values tended to be lower in dNES homozygous mice, neither AST nor ALT showed patterns indicative of hepatic injury. These findings support that the pathological phenotype of dNES mice is kidney-specific rather than systemic. Sample sizes were: WT (4 weeks), n = 14; WT (7 weeks), n = 16; dNES heterozygous (4 weeks), n = 13; dNES heterozygous (7 weeks), n = 7; dNES homozygous (4 weeks), n = 14; dNES homozygous (7 weeks), n = 6. Data are presented as mean ± SEM.
